# Transcriptional responses of acute glucose deprivation reveal a role for Snf12 and Spt20 in metabolic adaptation during stress

**DOI:** 10.64898/2026.08.17.745332

**Authors:** Justas Stanislovas, Kamilla Laidlaw, Katherine Paine, Daniel Ghete, Alastair Droop, Samantha Donninger, Sally James, Zoe Ingold, Amy Milburn, Chris MacDonald

## Abstract

The budding yeast *Saccharomyces cerevisiae* is a well-established model organism to study cellular stress response and underlying mechanistic regulation. Although glucose starvation fundamentally alters gene regulation and cell behaviour, inconsistent deprivation protocols often trigger gross morphological artefacts. These non-specific changes confound findings by activating pathways independently of true glucose-signalling mechanisms. Furthermore, a thorough transcriptomic profile of glucose starvation using non-confounding conditions remains lacking. Consequently, the precise transcriptional impact of losing key metabolic regulators that mediate adaptation to glucose starvation remains undefined. Here we have employed a refined glucose starvation protocol, utilising raffinose exchange, which shows induction of vast transcriptional stress response with minimal impact on cellular morphology confirmed by label-free imaging. Transcriptomic profiling revealed shifts in metabolic regulation, ATP turnover, and cell-to-cell communication as acute glucose deprivation driving cells towards oxidation-driven metabolism. Additionally, we characterise transcriptional alterations seen in deletion mutants of *SNF12* and *SPT20*, known regulators of cellular metabolism, showing previously unappreciated transcriptional conservation, in part mimicking glucose starvation response. Finally, we identified cargo and stress-specific expression related to both eisosome components and surface transporters that are critical for metabolic adaptation. Overall, this dataset provides a comprehensive transcriptomic resource for dissecting stress signalling and driving novel hypothesis generation.

## INTRODUCTION

Fluctuations in environmental nutrient availability present a universal challenge to cellular homeostasis, requiring organisms to rapidly calibrate their internal physiology to match external conditions for optimal cell behaviour. Among these metabolic cues, the deprivation of glucose serves as a primary, high-priority stress signal that forces an immediate adaptation from active cellular growth to energy conservation. For decades, the budding yeast *Saccharomyces cerevisiae* has served as a premier model organism for dissecting these fundamental responses. Yeast share much of the central machinery for glucose processing with other eukaryotic cells, including human cells. Insights gained from this simple microorganism continue to inform our understanding of human metabolic disease, with yeast serving as a particularly good model for rapidly growing mammalian cells, such as highly proliferative cancer cells (Diaz-Ruiz et al., 2011). Glucose acts as a master signalling molecule in yeast, orchestrating overlapping downstream signalling pathways that coordinate growth and development (Broach, 2012). Yeast has also been a useful model to study glucose starvation, as although during fermentative growth yeast prioritise glucose as a carbon source, but they have a series of alternative mechanisms to utilise alternative carbon sources (Kayikci and Nielsen, 2015). This has elucidated signal transduction pathways and been a powerful model for transcriptional regulators that respond to extracellular energy sources to employ different metabolic modes (DeRisi et al., 1997).

Classical genetic and biochemical research initially focused on the immediate metabolic adaptations to glucose starvation, detailing how yeast cells transition from rapid fermentation to respiratory metabolism during the diauxic shift, a process accompanied by the sweeping transcriptional repression of glycolytic enzymes and the activation of gluconeogenic pathways (Carlson, 1999). However, beyond modulation of central metabolism, this glucose starvation responses require a profound physical reorganisation of existing cellular features. This survival programme is governed by a network of evolutionarily conserved signalling hubs that sense fluctuating ATP levels and nutrient scarcity. Central among these is the Target of Rapamycin Complex 1 (TORC1) pathway, which is rapidly inactivated upon glucose withdrawal to halt energy-intensive anabolic processes like translation and ribosome biogenesis (Rødkær and Færgeman, 2014). Concurrently, nutrient-sensing kinases, most notably the Snf1 protein kinase (the yeast ortholog of mammalian AMP-activated protein kinase, or AMPK), are activated to orchestrate a global reprogramming of cellular physiology (Janapala et al., 2019). Crucially, these conserved signalling cascades do not merely alter gene expression; they converge directly on the plasma membrane (PM) and the endolysosomal trafficking machinery to execute rapid, post-translational remodelling of the cell surface (Becuwe et al., 2012; MacGurn et al., 2011). This internalisation does not merely clear the cell surface; the subsequent trafficking and cargo delivery to the vacuole facilitates the proteolytic degradation of these proteins, thereby generating a vital pool of recycled amino acids and nutrients to sustain cellular metabolism during prolonged starvation.

These conserved responses converge directly on plasma membrane (PM) and endolysosomal trafficking to execute physical remodelling. To minimise energy expenditure and recycle vital components, the cell rapidly alters its plasma membrane landscape, accelerating clathrin-mediated endocytosis (CME) to internalise non-essential nutrient transporters and routing them via endosomes to the vacuole for degradation. For example, glucose deprivation triggers translocation of the transcriptional repressor Mig1 from the nucleus, resulting in higher levels of the CME adaptors Yap1801 and Yap1802 to increase and promote degradation of surface proteins (Laidlaw et al., 2021). Concurrently, Mig1 also triggers elevated levels of Gpa2, which inhibits the Gpa1-PI3K mediated endosomal recycling of cargoes back to the PM (Laidlaw et al., 2022). Furthermore, glucose starvation causes specialised membrane microdomains like eisosomes sequester vital cargo to ensure rapid post-starvation recovery (Laidlaw et al., 2021; Paine et al., 2023) and expanded organelle contact sites, thought to facilitate vacuolar maintenance for changing conditions (Hugenroth et al., 2025). Consequently, analysing glucose starvation in yeast provides an elegant, physiologically relevant paradigm to dissect how universally conserved metabolic sensors directly cross-talk with vesicle sorting and organelle dynamics.

However, standard methodologies for studying glucose starvation in *Saccharomyces cerevisiae* often suffer from confounding experimental artefacts. Transferring yeast directly to sugar-free/glucose-depleted media drastically alters external osmolarity. This triggers the High-Osmolarity Glycerol (HOG) pathway, inducing rapid changes in turgor pressure (Hohmann, 2002). Furthermore, spheroplast allows for efficient RNA extractions but this process can cause large adaptions via cell wall integrity pathways, alters mechanosensitive ion channels, and disrupts spatial organelle organisation. As such, performing this process before applying stress conditions (Su et al., 2023) will grossly affect transcriptomic readings. Finally, early transcriptomic studies that are still commonly used for reference relied on low-resolution microarray datasets that lack the sensitivity, dynamic range, and resolution to capture subtle non-coding RNAs, transcript isoforms, or subtle shifts in low-abundance membrane-bound protein machinery.

In this study we employ an alternative carbon source raffinose in the media to maintain osmotic balance, while depriving cells specifically of glucose. Label free imaging shows this induces less physiological shock to cells whilst effectively cancelling glucose metabolism pathways, allowing pathways specific to glucose metabolism to be disentangled from other stress responses. We use our optimised starvation protocol to perform a series of RNA sequencing (RNAseq) experiments to profile intact cells, comparing glucose to raffinose media but also focussing on some global transcriptional regulators related to metabolism (Spt20and Snf12), to generate a datasets that bypasses osmotic shock and mechanical artefacts. This isolates the true glucose-starvation transcriptional program without confounding stress responses. Analysis focuses on key changes and discusses how eisosome related factors that protect against stress are regulated at the transcriptional level.

## RESULTS

### Raffinose exchange induces glucose starvation with reduced physiological effects

We have previously employed holotomography to study yeast cell parameters, which is a powerful label free imaging technique that can be performed in synthetic media but also auto fluorescent rich yeast media that is typically incompatible with fluorescence microscopy (Xelhuantzi et al., 2024).To build on this, we employed a high throughput holotomography instrument coupled to automated segmentation and analysis pipeline. Using this approach, we could capture the same fields of view before and after starvation of glucose. We used HT-X1 to image large pools of yeast cells and the TomoAnalysis platform was used to segment and quantify yeast cells in 3D (**Figure 1A**). These experiments show that physical parameters, such as reduced area and volume, were much more affected by shifting to glucose free media compared to media supplemented with raffinose (**Figure 1B**). We have previously shown that cells exposed to glucose for 6 hours behave as if they are starving for glucose, with high levels of surface localised glucose transporters retained at the surface (Laidlaw et al., 2021). However, the immediate effects in the shift are obvious, with a stall in growth of raffinose treated cells obvious following 30 minutes treatment (**Figure 1C**).

**Figure 1:**
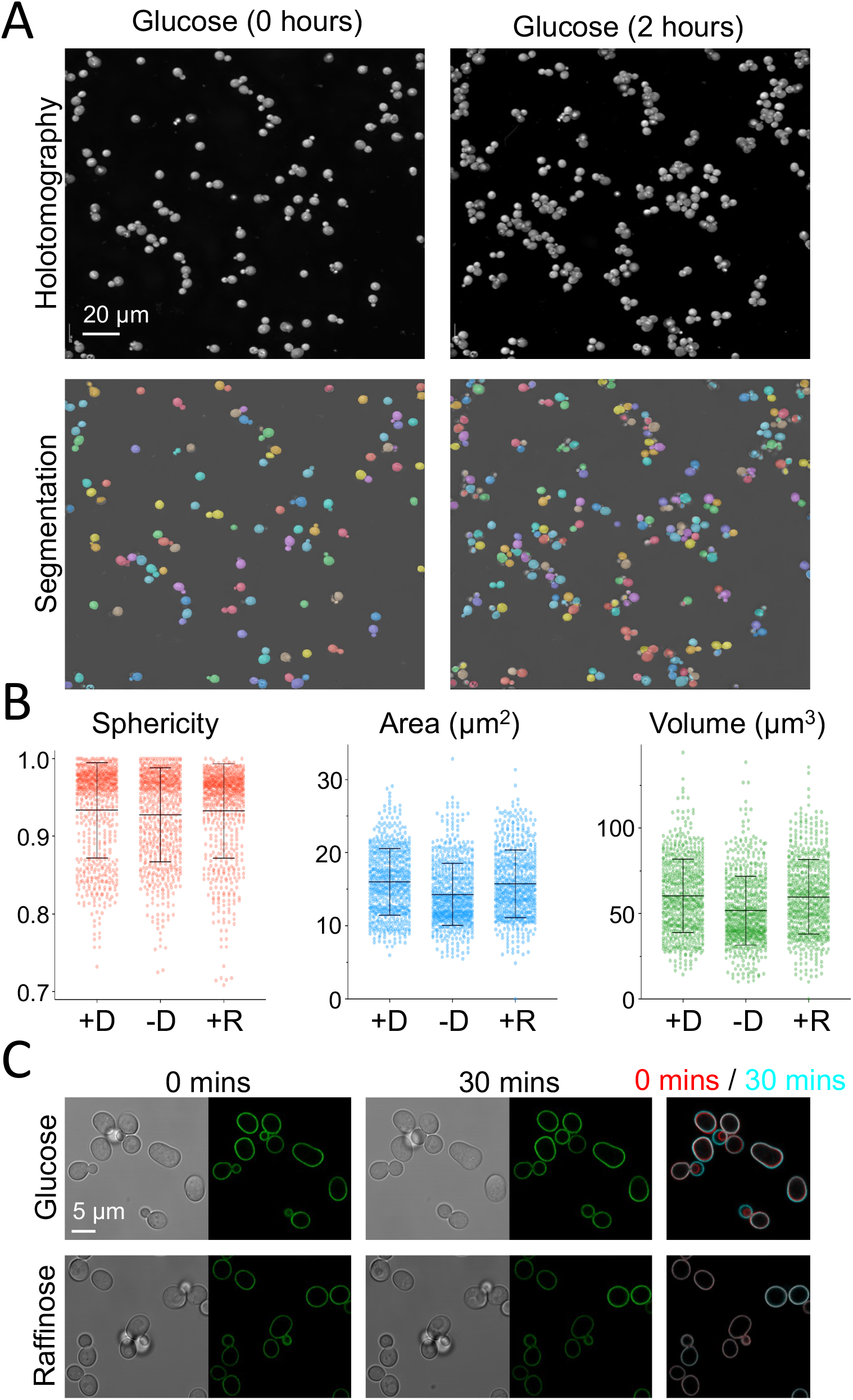
Glucose starvation mediated by raffinose exchange results in physiological adaptation. **A)** Time lapse holotomography was used to image yeast cells before (0 hours) and after (2 hours) of raffinose exchanged media. Fields of view were segmented using the Tomocube analysis software, successfully identifying individual cells for downstream analysis. **B)** Analysis of yeast cells from holotomograms, showing cells exposed to glucose (+D), glucose starvation in a water-based media (-D) and following exchange with raffinose media (+R). **C)** Time lapse imaging was used to exchange mid-log phase yeast cells from glucose containing media to a raffinose equivalent. Cell growth was compared at 0 minutes (red) and after 30 minutes exchange (cyan).

### Transcriptional changes induced following raffinose exchange

Based on our previous experiments, we set out to assess transcriptional responses using bulk RNAseq to compare cells grown in glucose with cells exposed to raffinose for 6 hours (**Figure 2A**). For this analysis we also included two key regulators of gene expression in yeast: the Snf12 member of the SWI/SNF chromatin remodelling complex and Spt20, a subunit of the SAGA transcriptional regulatory complex (**Figure 2B**). We have generated Spt20 and Snf12 genomic knockouts (*spt20Δ* and *snf12Δ*), and in parallel to glucose starvation, assess the role of Spt20 and Snf12 in transcriptional regulation. Using mRNA-enrichment we have sequenced 10 samples of WT cells (glucose grown), 5 samples that have undergone raffinose exchange, and 5 samples of each *spt20Δ* and *snf12Δ* mutants (**Figure 2C**). Principal Component Analysis (PCA) shows transcriptional variance indicating high level of conservation within replicates of each condition, and clear transcriptional differences between WT, glucose grown, and raffinose exchanged media samples (**Figure 2C**). Interestingly we see some transcriptional similarity between *spt20Δ* and *snf12Δ* mutants, as they cluster together. In agreement with PCA, the top 500 most variable genes indicate robust transcriptional differences and transcriptional reproducibility of replicates (**Figure 2D**). Collectively this indicates vast, transcriptome-level changes in gene transcription upon loss of glucose and Snf12 and Spt20-mediated transcriptional regulation.

**Figure 2:**
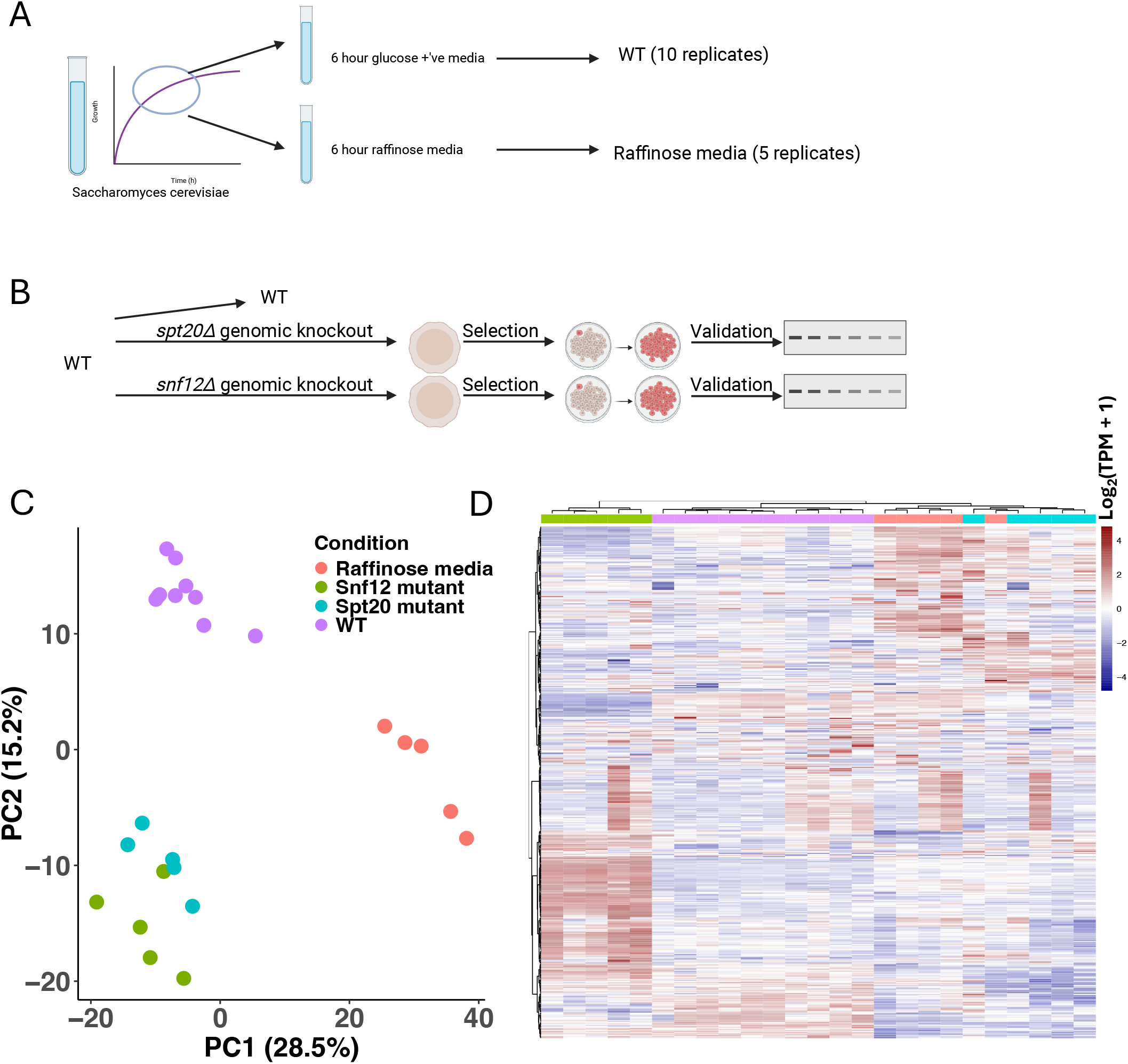
Transcriptional profiling of raffinose-stressed and *spt20Δ* and *snf12Δ* mutant strains. **A)** Schematic depicting experimental design generating WT and raffinose media-exchanged samples. Following growing WT cells to early log phase, cells were then brought up for additional 6 hours either in glucose rich media or raffinose media lacking glucose. **B)** Schematic depicting generation of *spt20Δ* and *spt20Δ* mutant strains. **C)** Principal Component Analysis (PCA) of the top 500 most variable genes of raffinose-exchanged samples, *snf12Δ* mutant, *spt20Δ* mutant and WT samples. PC1 and PC2 denotes principal component 1 and 2, respectively. **D)** Heatmap of log2 + 1 normalized transcripts per million (TPM) of the top 500 most variable genes. Refer to the legend in panel C for column annotation in panel D. **B** and **C** were created in BioRender. Stanislovas, J. (https://BioRender.com/ed8wmc4) is licensed under CC BY 4.0

We first assessed transcriptional differences between WT cells grown to log phase in glucose replete media or following acute (6 hrs) raffinose exchange using differential gene expression (DGE) analysis, yielding a vast gene set of significantly differentially expressed genes, designated to be absolute log2 fold change (log2FC) of 1 (upregulated at > log2FC of 1, downregulated at < log2FC of -1), at the p-adjusted value of less than 0.1 (**Figure 3A**). Out of a total of 652 differentially expressed genes at the chosen cut-off, most significantly upregulated and downregulated genes indicate transcriptional metabolic rewiring (**Figure 3B**), through, for example, upregulation of amino acid transporters, kinases involved in fatty acid synthesis, and utilisation of glycerol. The most upregulated gene in raffinose, *JEN1*, is a known transmembrane transporter of lactate, pyruvate and acetate, normally transcriptionally repressed in the presence of glucose (Andrade et al., 2005; Chambers et al., 2004; Paiva et al., 2009). Of note, we also detect a plethora of significantly differentially expressed unannotated protein-encoding transcripts, suggesting potential novel regulators of adaptation to stress response (**Figure 3C**). At the pathway level, gene set enrichment analysis (GSEA) indicates majority of gene ontology (GO) terms across all significant biological processes (BP) to be associated with oxidative metabolism, such as electron transport chain (ETC), adenosine triphosphate (ATP) generation, and other oxidation-dependent processes (**Figure 3D**). Indeed, the top enriched GO BP terms, molecular functions (MF) and cellular compartments (CC), shows upregulation of mitochondrial translation and ETC, indicating a shift towards oxidative metabolism in the exchange of glucose (**Figure 3E**). The second most common category of enriched BP terms, sugar metabolism, including terms such as hexose and maltose transport, indicates cellular tendency to replace glucose with an alternative carbohydrate source for energy derivation in the absence of glucose (**Figure 3D**). In agreement with our live cell imaging data, indicating mild cellular response to acute stress during raffinose exchange (**Figure 1**), we detect mild suppression of endoplasmic reticulum (ER) activity and cell wall organisation (**Figure 3D - E**), but no other forms of cellular stress, whilst post-translational modifications and protein localisation processes are also suppressed in raffinose. This confirms that raffinose is a mild physiological stressor without gross cellular changes, whilst inducing profound transcriptional changes spanning a range metabolic and energetic processes governing life, highlighting its usefulness in studies of stress response signalling.

**Figure 3:**
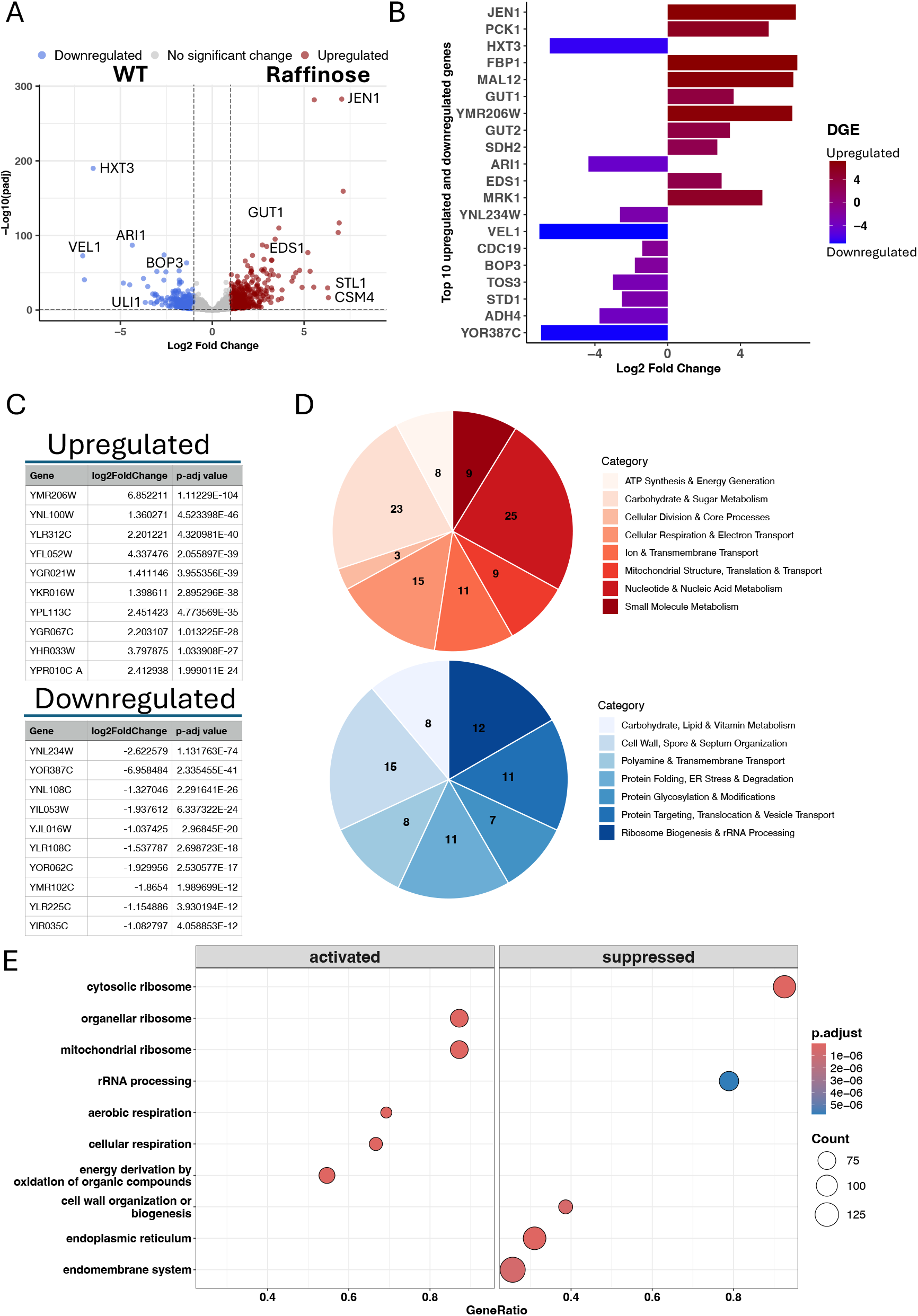
Glucose starvation induces transcriptional changes indicating metabolic rewiring and cellular adaptation. **A)** Volcano plot showing differential gene expression (DGE), with significantly differentially expressed genes considered to be at absolute log2 Fold Change of 1 (vertical dashed lines indicating cut-offs of -1 for downregulated and 1 for upregulated) and –adjusted value of 0.1 (horizontal dashed line). **B)** Top 10 most significantly upregulated and downregulated genes in raffinose vs WT identified in DGE, with genes being listed in the order of most significant (*JEN1*) to least significant (*YOR387C*). C) Top 10 novel transcripts identified to be most significantly differentially expressed in DGE. **D)** Gene set enrichment analysis (GSEA) of Gene Ontology (GO) identified biological processes (BP) terms that are enriched (top pie chart, red colour palette) and suppressed (bottom pie chart, blue colour palette) in raffinose compared to WT based on the DGE list at p-adjusted value of <0.1. The numbers indicate the number of terms that belong to that category. **E)** Top 10 GO terms activated and suppressed in raffinose compared to WT.

### Conserved Spt20 / Snf12 transcriptional control of metabolic processes

To assess if transcriptional alterations in response to mild cellular stress, validating our approach, are similar to changes induced by the loss of two key transcriptional regulators with known effects on metabolism, Spt20 and Snf12, we have performed DGE between each null mutant and WT strain, yielding a vast list of differentially expressed genes (**Figure 4A - B**). At the cut-off of absolute log2FC of 1 and p-adjusted value < 0.1, a total of 405 and 345 genes were significantly altered in expression in Snf12, and Spt20 mutants compared to WT, respectively. Some of the top differentially expressed genes in *spt20Δ* and *snf12Δ* mutants vs WT include regulators of metabolism, cell growth, and cell communication (**Figure 4A - B**), in line with their role in global control of gene transcription (Holstege et al., 1998). DGE list (all genes at p-adjusted values < 0.1) yields GSEA where we observe similarity in the top enriched and suppressed GO terms (**Figure 4C - D**), notably activation of sporulation related terms and suppression of cytoplasmic translation, akin to downregulation of translation in raffinose, indicating transcriptional control of similar biological functions and the role of SWI/SNF chromatin remodelling complex and the SAGA complex in governing protein translation. Given the similarity of altered biological processes, we have analysed the overlap in the DGE list between *SPT20* and *SNF12* deletion mutants vs WT, revealing a substantial number of genes that are concordantly upregulated or downregulated in both mutants compared to WT (**Figure 4E - F**). Whilst some of this may still be stochastic transcriptional signal, utilising STRING analysis searching for interactions at medium confidence (0.4) depicted by the edges connecting the nodes, we observe a number of overlapping genes that belong to the same functional group, such as genes involved in thiamine synthesis, non-glucose sugar metabolism and transport, upregulated in both mutants (**Figure 4G**), and genes involved in processes such as biotin synthesis and mating were found downregulated in both mutants (**Figure 4H**). Collectively, these observations suggest that 1) loss of either regulator with no other changes induce vast transcriptional changes, 2) loss of Spt20 and Snf12 induce cellular stress similar to raffinose exchange highlighting their role in control of metabolic regulation, and 3) Spt20 and Snf12 may show previously underappreciated transcriptional conservation in regulation of downstream signalling.

**Figure 4:**
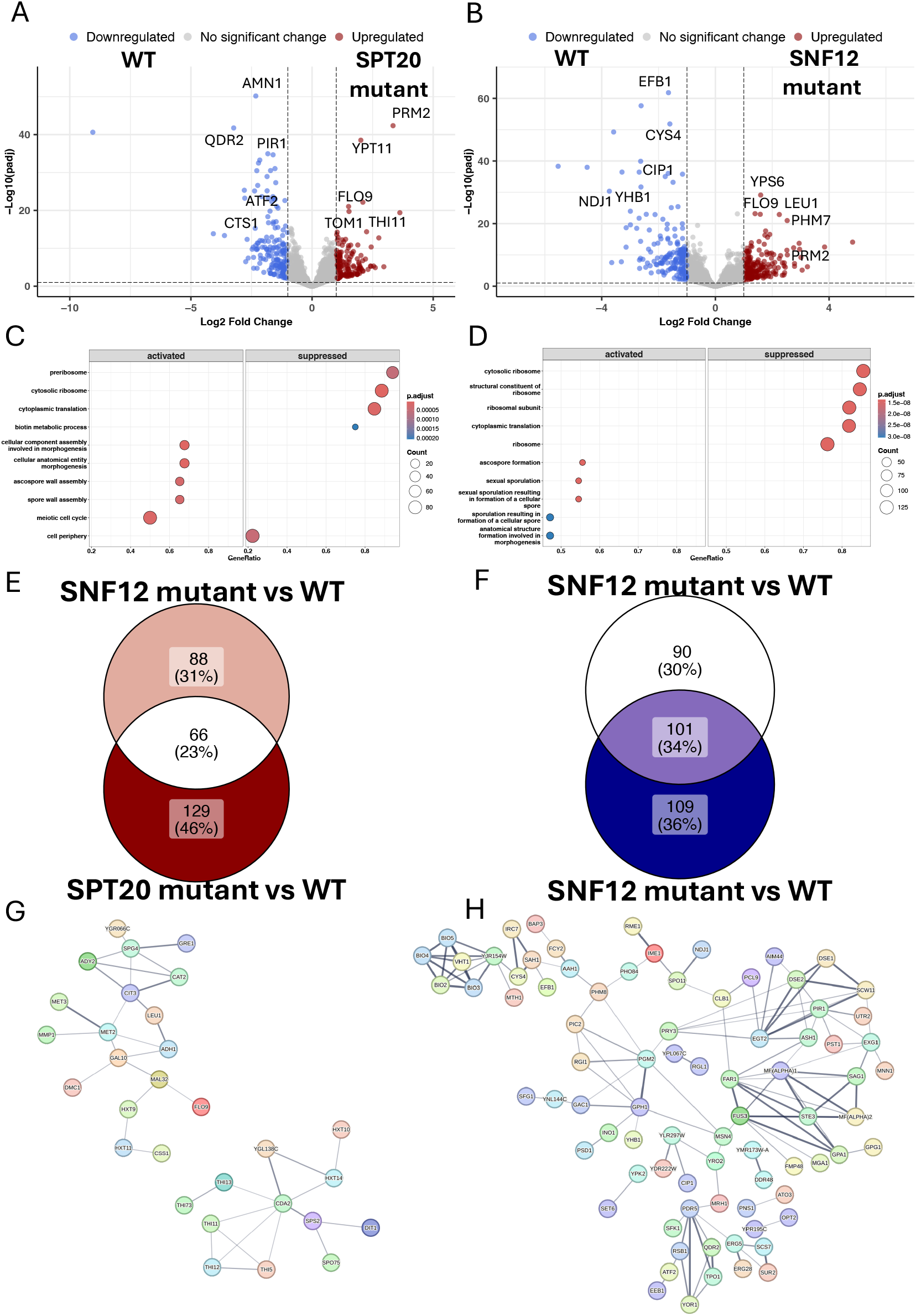
*spt20Δ* and *snf12Δ* mutants show conservation in their transcription profiles. **A)** Volcano plots showing DGE between *spt20Δ* and WT and **B)** *snf12Δ* mutant and WT, with significantly differentially expressed genes considered to be at absolute log2 Fold Change of 1 (vertical dashed lines indicating cut-offs of -1 for downregulated and > 1 for upregulated) and –adjusted value of 0.1 (horizontal dashed line). **C)** Top 10 GO terms (across biological processes, molecular functions and cellular compartments) activated and suppressed in *spt20Δ* vs WT, and **D)** in *snf12Δ* vs WT based on the DGE list at p-adjusted value of <0.1. **E)** and **F)** Venn diagrams depicting the number and percentage of significantly uniquely and commonly upregulated and downregulated genes, respectively, in *snf12Δ* vs WT and *spt20Δ* vs WT DGE analysis. **G)** and **H)** STRING network analysis showing potential interactions identified at 0.4 confidence score limit (thickness of the edge denotes how strong the confidence) across all possible interaction sources, textmining, experiments, databases, co-expression, neighbourhood, gene fusion, and co-occurrence, in genes commonly upregulated and downregulated in *spt20Δ* and *snf12Δ* mutant vs WT, respectively. Gene list depicted excludes genes that had no identified interactions.

### Eisosomal specific responses to glucose starvation are transcriptionally different to loss of Snf12 and Spt20

Our transcriptomic characterisation indicates large scale transcriptional rewiring upon raffinose exchange and deletion of *SNF12* and *SPT20*, conditions inducing stress response changes without gross cellular defects. As eisosomes are quintessential to rapid post starvation recovery given their role in sequestration of metabolic cargo (Gournas et al., 2018; Laidlaw et al., 2021; Moharir et al., 2018), and our data indicate acute starvation response and transcriptional metabolic rewiring, we analysed expression of key eisosome genes, genes regulated by eisosomes, and eisosome-associated transporters across our transcriptomic dataset (**Figure 5A**). As expected, in WT cells grown in glucose, eisosome genes show generally low expression levels, consistent across all biological replicates. Whilst showing greater variance between samples, cells exposed to raffinose show clearly different transcriptional profile of eisosome expression with elevated gene expression of many eisosome genes (**Figure 5A**). Indeed, several eisosome genes and surface transporters are significantly upregulated in raffinose vs WT as seen in DGE, underlying their role in metabolic adaptation through regulation of surface cargo trafficking (**Figure 5B**). The top upregulated eisosome gene, *MDG1*, component of the core eisosome Pil1 complex, to the best of our knowledge has not been previously shown to be transcriptionally regulated in response to acute glucose starvation (**Figure 5C**). The most upregulated surface transporter in response to raffinose exchange. Also, MMP1, encoding S-methylmethionine transporter (**Figure 5C**), suggests increased need for sulphur compounds, likely to compensate for oxidative stress which is induced by the loss of glucose. In parallel, some eisosome components and surface transporters are downregulated in response to glucose starvation (**Figure 5B, 5D**), suggesting likely cargo-selective transcriptional alterations. Although raffinose exchange induces modulation of Pil1 stoichiometry at single molecule resolution (Laidlaw et al., 2021), and triggers rapid Pil1 dephosphorylation (Paine et al., 2023), we have previously observed only modest and acute transcriptional changes in *PIL1* that plateau after <2 hours, consistent with our findings of no change at 6 hours (**Figure 5B**). In contrast, *PIL1* expression shows robust transcriptional downregulation in *spt20Δ* mutants but not *snf12*Δ mutant cells, which is corroborated at the protein expression level (**Figure 5E**). This may suggest specific and acute regulatory control of eisosomes in response to drastic metabolic stress conditions.

**Figure 5:**
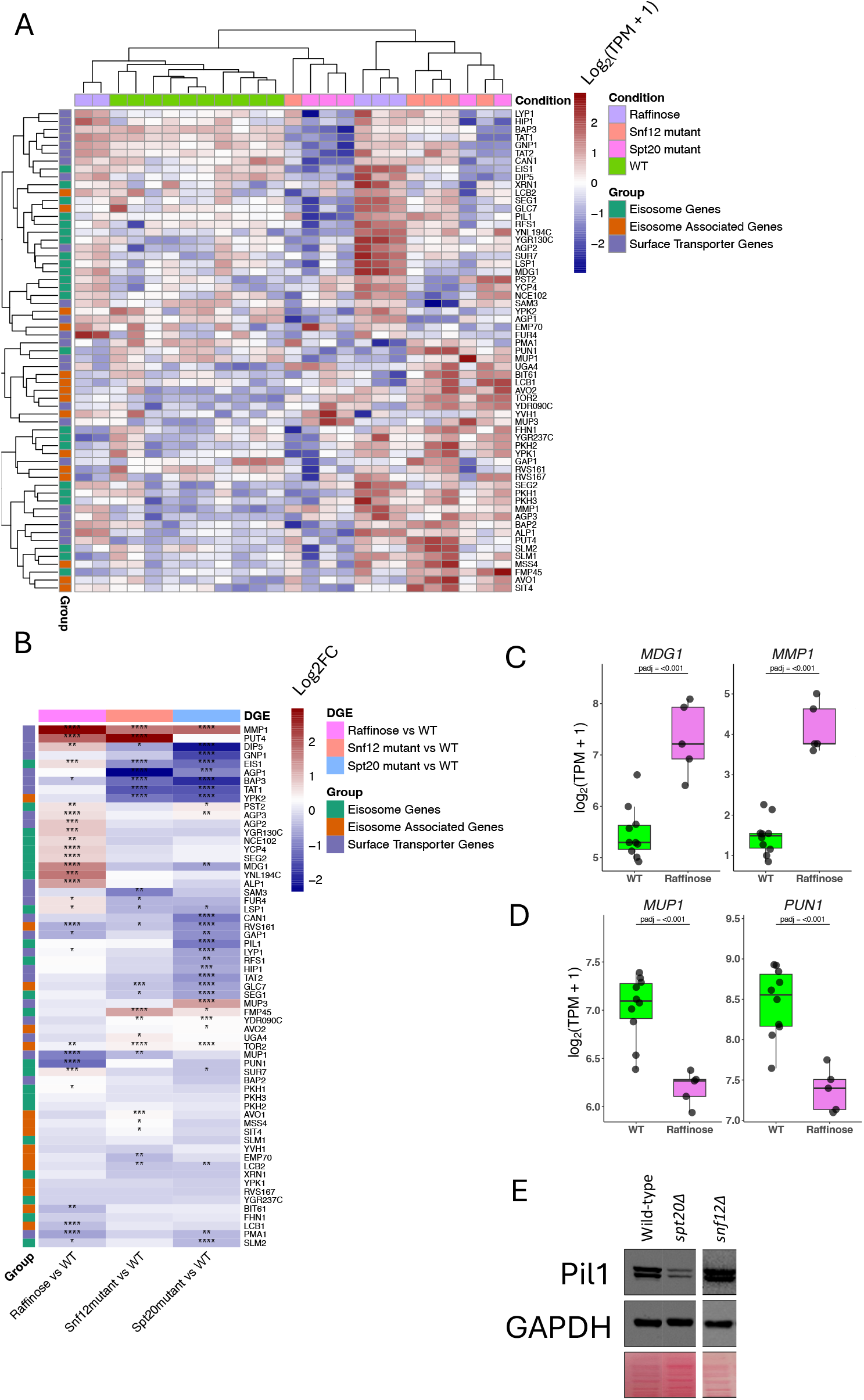
Glucose starvation promotes expression of select eisosome genes and surface transporters different to *snf12Δ* and *spt20Δ* mutants. A) Heatmap of counts (log2(TPM+1)) of select, known eisosome genes, eisosome associated genes and surface transporter genes across all replicates of experimental conditions. **B)** Heatmap of log2FC values from DGE analysis between raffinose vs WT, *snf12Δ* mutant vs WT, and *spt20Δ* mutant vs WT. The scale depicts log2FC values directly from Deseq2 analysis, with the asterisks denoting p-adjusted value, with *p-adj < 0.05, **p-adj < 0.01, ***p-adj < 0.0001, ****p-adj near 0. **C)** Boxplots of the most significantly upregulated eisosome and surface transporter gene in raffinose vs WT, and **D)** boxplot of the most significantly downregulated eisosome and surface transporter gene in raffinose vs WT. p-adjusted value is taken directly from DGE analysis using Deseq2. **E)** Western blots of Pil1 levels in WT, *spt20Δ* mutant and *snf12Δ* mutant.

Indeed, in contrast to exposure to raffinose, *snf12Δ* and *spt20Δ* mutants show largely conserved downregulation of eisosome genes, their regulated genes and associated surface transporters (**Figure 5B**), indicating that despite Snf12 and Spt20 regulation of metabolic processes highlighted earlier, loss of Snf12 and Spt20 does not enhance surface cargo internalisation and trafficking. Interestingly, *MMP1* is also upregulated in both *snf12Δ* and *spt20Δ* mutant conditions, suggesting some response to oxidative stress known to be regulated by SWI/SNF1 complex which Snf12 is a part of, and SAGA complex, including Spt20. On the other hand, differences in upregulation/downregulation state of transcriptional change in each condition indicates that whilst there is a basal level of oxidative stress seen across raffinose exchange and loss of and Snf12 and Spt20, the gene-level regulation is not conserved. For example, whilst *DIP5*, dicarboxylic acid permease, shows upregulation in raffinose, it is downregulated in *spt20Δ* mutant cells. Similarly, despite oxidative stress in the absence of glucose, *MUP1*, a methionine transporter, is only upregulated in *spt20Δ* mutant, whilst raffinose conditions induce *MUP1* downregulation (**Figure 5B - C**). Collectively these findings suggest intricate transcriptional network of regulation of surface cargo internalisation and trafficking, likely in a substrate-specific manner, in different stress response conditions in part fuelled by oxidative insult.

## DISCUSSION

Acute glucose starvation triggers rapid, wide-ranging shifts in yeast cell biology. However, conventional protocols often introduce starvation by stripping all sugars from the growth media, unintentionally inducing severe osmotic shock. These approaches confound true nutrient-deprivation signalling with hyperosmotic stress, and has complicated the interpretation of historical datasets, leaving key adaptive mechanisms incompletely understood. Here we have employed a refined glucose-starvation protocol that avoids gross, artefactual cellular changes in osmotic potential to study transcriptional response to acute loss of glucose as a quintessential metabolic stress in *S. cerevisiae*. We previously demonstrated that transferring cells to raffinose-containing media avoids the widespread downregulation of distinct high-affinity hexose transporters (Hxt6 and Hxt7 tagged with GFP) typically observed during acute glucose deprivation (Laidlaw et al., 2021). Unlike monosaccharides, raffinose requires extracellular enzymatic cleavage prior to cellular import and utilisation (Fuente and Sols, 1962). This dependency creates a controlled, low-flux carbon regime rather than a sudden metabolic collapse. Consequently, raffinose treatment induces far milder morphological alterations (**Figure 1**), bypassing the indirect structural artefacts, which may confound true biological response, such as the disorganisation of eisosome microdomains and impaired endocytic trafficking, that can be triggered in response to cell size and membrane tension changes (Riggi et al., 2019, 2018; Wu et al., 2017). We also note that using label-free imaging allows rapid and accurate cell segmentation of even densely populated regions, in addition being efficient in rich yeast media, that is suboptimal for fluorescence microscopy (Xelhuantzi et al., 2024). We therefore propose that raffinose treatment provides a more physiologically faithful model of glucose starvation than total carbon deprivation, due to maintaining a more stable osmotic balance. By establishing this refined baseline, the downstream transcriptional changes documented in this study offer greater physiological relevance and functional specificity. Using 3D holotomography to capture these structural shifts offers clear analytical advantages. This technique enables high-throughput, label-free 3D imaging of living cells, combining rapid image acquisition with efficient automated segmentation.

Indeed, bulk RNAseq-generated transcriptomic datasets of wild-type cells grown in glucose versus raffinose shows few signs of morphological cellular stress, but robust and vast transcriptional rewiring spanning ATP turnover, sugar and amino acid metabolism and cell-to-cell communication. In agreement with the retention of high-affinity hexose transporters at the cell surface 6 hours after raffinose exchange (Laidlaw et al., 2021), we also observed a stark downregulation of *HXT3*, consistent with the cell actively shutting down low-affinity transport machinery while maintaining high-affinity scavenging capabilities (**Figure 3**). Concurrently, this analyses also shows this glucose-limited state triggered a massive upregulation of *JEN1*, marking a broader transition towards alternative carbon utilization pathways once glucose repression is relieved aligning with known activation (Becuwe et al., 2012; Becuwe and Léon, 2014; Chambers et al., 2004). Whilst we report large-scale transcriptional characterization of changes induced by glucose starvation, the dataset presented is vast in both the number and diversity of pathway alterations, as well novel gene regulators, which can help generate detailed mechanistic hypotheses. Moreover, we further strengthen the richness of this transcriptomic dataset by including deletion mutants of Spt20 and Snf12, fundamentally involved in broad metabolic regulation (Holstege et al., 1998; Lee et al., 2000), including glucose metabolism. Collectively this creates a unique set of transcriptomic conditions that we integrate into a comprehensive analysis of transcriptional stress response.

Given the fundamental role of eisosomes in internalising and trafficking cell-surface cargoes, transcriptomic assessment of their expression reinforces their importance in adapting to glucose starvation. Although eisosomes protect resident proteins from endocytosis, retaining nutrient transporters within these domains provides a distinct physiological advantage during recovery from nutrient deprivation (Appadurai et al., 2020; Gournas et al., 2018; Laidlaw et al., 2021; Moharir et al., 2018; Paine et al., 2023) starvation conditions like glucose deprivation. In addition to these glucose-regulated responses, our data highlight a broader role for the underlying transcriptional machinery. By evaluating *spt20Δ* and *snf12Δ* mutants, we uncover a complex regulatory network that not only controls eisosome expression, likely to optimise transporter retention during starvation, but also coordinates the transcriptional repression of numerous nutrient transporters (**Figure 5**). These findings provide conceptual validation for our model and offer a framework for future studies to dissect why specific eisosomal components are selectively upregulated during glucose deprivation to facilitate post-starvation recovery.

We also report novel eisosome-like genes to be upregulated in glucose starvation, which have previously not been identified as candidates of cargo trafficking. Further, in light of lack of upregulation of *PIL1* shown previously and confirmed in this study, yet transcriptional upregulation of some of Pil1 complex components, such as *MDG1* in raffinose conditions, may suggest transporter regulation is mediated by less well understood factors like Mdg1, as opposed to Pil1 that serves a core eisosome biogenesis function (Walther et al., 2007, 2006). Recently Mdg1 has been shown to transcriptionally regulate activity of a range of metabolic enzymes and regulators of the cell cycle (Wan et al., 2026), implying a link to metabolism mediated through eisosomes and downstream responses. Although Spt20 and Snf12 are associated with broad transcriptional changes (Holstege et al., 1998; Lee et al., 2000), they are also inherently linked to glucose metabolism: with Spt20 SAGA complex required for the transcriptional activation of stress-induced and nutrient-responsive genes. (Grant et al., 1997; Roberts and Winston, 1996) and Snf12 being first identified through sucrose utilisation mechanisms (Carlson et al., 1981). Investigating *spt20Δ* and *snf12Δ* mutants alongside glucose-starved cells provides a direct probe into how chromatin co-activators coordinate global metabolic adaptation. Because Spt20 and Snf12 serve as key structural subunits of the SAGA acetyltransferase and SWI/SNF nucleosome remodelling complexes respectively, transcriptomic profiling of these deletion strains pinpoints the baseline gene networks reliant on epigenetic chromatin architecture. Assessing these mutant profiles in parallel with the transcriptional shifts induced by glucose starvation reveals whether nutrient-deprivation signals recruit SAGA and SWI/SNF to derepress metabolic genes, or if alternative pathways compensate when these major chromatin remodelling complexes are compromised. Indeed, we see previously unappreciated overlap in transcriptional profiles of *snf12Δ* and *spt20Δ* mutants, suggesting a level of conservation in their regulation of gene expression, converging on metabolic adaptation, akin to seen in glucose starvation. Whilst, for example, SAGA and SWI/SNF complex have both been shown to regulate cell wall integrity in a cooperative manner (Sanz et al., 2016, 2012), less is known about their common regulation of thiamine synthesis of cell-to-cell communication. Moreover, this rich dataset provides a comprehensive transcriptomic dataset resource to generate pathway or gene-level hypotheses to help mechanistically untangle transcriptional regulation of stress responses induced by different stimuli. Similarly, the transcriptional networks that control other membrane trafficking pathways and the expression of specific transporters help understand the key factors in adapting to glucose starvation. This work will therefore have implications, for understanding other work in yeast, and as many of these factors are conserved, also impact other eukaryotic studies.

## METHODS

### Reagents

Wild-type yeast used were BY4742 strain (Brachmann et al., 1998) , which also served as the parental strain for the *spt20Δ::G418* and *snf12Δ::G418* mutants (Giaever et al., 2002). Pil1 anti-rabbit polyclonal antibodies were a kind gift from Professor Tobi Walther (Harvard Medical School) and used as described in their original article (Walther et al., 2007). α-GAPDH anti-mouse monoclonal antibodies (clone 6C5) were also used (Santa Cruz, CA, USA). For details, see immunoblotting section below.

### Cell culture

Prior to transcriptomic analyses, individual colonies were isolated for wild-type (on YPD), and both *spt20Δ* and *snf12Δ* mutants (on G418). 10 colonies from wild-type, and 5 from each mutant were phenotyped using auxotrophic markers and for the mutants genotyped to ensure *SPT20* and *SNF12* genes were deleted. Cell were then grown in serial dilution at 30°C, with 8x 10-ml cultures (diluted 2-fold) grown overnight in synthetic complete media (2% glucose, with full complement of yeast nitrogen base and amino acids and base mixtures (Formedium, UK) so that log phase cultured could be harvested the next morning. For raffinose exchange media., wild-type cells were cultured exactly the same, but in the morning were washed 3x with SC media containing 2% raffinose instead of glucose), and then cultured in raffinose media for 6 hours at 30°C. Following growth, 20 x OD_600_ units of yeast culture were harvested, washed once and pellets flash frozen in liquid nitrogen and stored at -70 °C until ready for RNA extraction.

### RNA extraction & RNA-Seq library preparation

RNA extraction and RNA-Seq library preparation was performed by the Genomics lab of the Bioscience Technology Facility, University of York. Spheroplasting was performed immediately prior to RNA extraction by resuspending cell pellets in Y1 buffer (1 M sorbitol, 0.1 M EDTA, 0.2 % b-mercaptoethanol, 25 U/ml zymolyase) and incubating for 5 minutes at room temperature with gentle agitation. RNA was then extracted using Qiagen RNeasy RNA mini kits, according to the manufacturer’s instruction, and including an on-column DNase treatment step. RNA quantity and quality was assessed using the Agilent Bioanalyzer, and 1 μg high quality RNA was taken forward into library preparation. PolyA mRNA was isolated using the NEBNext polyA mRNA magnetic isolation kit, and sequencing libraries were prepared using the NEBNext RNA Ultra II Directional library preparation kit, with unique dual indices (New England Biolabs). The resultant sequencing libraries were pooled at equimolar ratios and subject to paired end 150 base sequencing on an Illumina NovaSeq X, to an average depth of approximately 10-15 million read pairs per sample.

### RNA-Seq analysis and Data & Code availability

RNAseq pair-end sample sequences (FASTQ files) were aligned and transcripts quantified using standard Salmon pipeline. DESeq2 R package (version 1.50.2) was then used to perform differential gene expression analysi, with the chosen cut-offs discussed in the results. All downstream bioinformatic analyses were performed using R version 4.5.0 (2025-04-11) in R studio. Gene counts and code used to generated analysis will be available upon requestion upon publication. . STRING web interface (string-db.org, version 12.0) was used to analyse gene-to-gene interactions at the cut-offs discussed in the results.

### Holotomography

Yeast were cultured to log phase and then prepared for holotomography at room temperature, using a 3D quantitative phase imaging (QPI) using an HT-2H instrument (Tomocube Inc), employing Mach–Zehnder interferometry and a digital micromirror device. 48x 2D holographic images acquisitions were collected, controlled by a digital micromirror device (DMD). These images were then used to construct a 3D tomogram based on refractive index, as described previously (Xelhuantzi et al., 2024). The diffracted beams from the sample were collected using a high numerical aperture objective lens (NA=1.2, UPLSAP 60XW, Olympus) and recorded using a complementary sCMOS image sensor (Blackfly S BFS-U3-28S5 M, FLIR Systems Inc). The data was imaged and visualized using TomoStudio (Tomocube Inc., Korea).

### Confocal microscopy

Yeast cells expressing GFP-Pmp3 (Weill et al., 2018; Yofe et al., 2016) as a marker for the yeast plasma membrane (Cruz-Xelhuantzi et al., 2026) were grown to mid-log (OD_600_ ≤ 0.5) phase in SC medium containing glucose then adhered to 35-mm glass-bottom coverslip dishes (Ibidi GmbH) coated with 1 mg/ml concanavalin A (Sigma-Aldrich) in water for 5 min prior to three washing steps. Bright field and fluorescence images of adhered cells were captured on a Zeiss laser scanning confocal instrument (Zeiss LSM980), using an Airyscan 2 detector and a 63×/1.4 objective lens. GFP was excited using a 488 nm laser and emission collected from 495 to 500 nm. For time lapse, following initial; images, sterile media exchanges with Sc media contaioning 2% raffinose (no glucose) were performed using 50-ml syringes through tubing fused to the lid of the 35-mm dishes. To exchange media, the dish was filled and emptied 3x with raffinose media, before a final incubation in media and time lapse images of the same field of view captured. All images were processed using Zeiss Zen software and modified (e.g. coloured) using ImageJ software (NIH).

### Immunoblotting

Strains were grown to mid-log phase, equal cell numbers estimated by optical density (OD_600_) prior to harvesting. Cell pellets were treated to 0.2 N NaOH for 5 min prior to resuspension in lysis buffer (8 M urea, 10% glycerol, 50 mM Tris-HCl pH 6.8, 5% SDS, 0.1% Bromophenol Blue and 10% 2-mercaptoethanol). Lysates were loaded on to 10% acrylamide SDS-PAGE gels and electrophoresis performed at constant voltage 200 mV). Resolved proteins were transferred to a nitrocellulose membrane using the iBlot2 dry transfer system (Invitrogen). Membranes were probed with antibodies, either 1/1000 (anti-Pil1) or 1/5000 (anti-GAPDH), overnight at 4°C, prior to washing in PBST (1x PBS, 0.2% tween), incubationg with IgG anti-rabbit (for Pil1) and anti-mouse (for GAPDH) secondary antibodies conjugated to HRP (Thermo Fisher Scientific). Immunoblots were visualised using enhanced chemiluminescence (ECL) Super Signal Pico Plus (Thermo Fisher Scientific) and captured using a ChemiDoc Imager (Bio-Rad).

## ACKNOWLEDGMENTS

We would like to thank staff at the York Bioscience Technology Facility for technical assistance. This research was supported by a Sir Henry Dale Research Fellowship from the Wellcome Trust and the Royal Society 204636/Z/16/Z (CM).

